# Greater capacity to exploit warming temperatures in northern populations of European beech is partly driven by delayed leaf senescence

**DOI:** 10.1101/707307

**Authors:** Homero Gárate-Escamilla, Craig C. Brelsford, Arndt Hampe, T. Matthew Robson, Marta Benito Garzón

**Author notes:** Corresponding author, BIOGECO UMR 1202, INRA - Université de Bordeaux, Bat B2, Allée Geoffroy-St-Hilaire, CS50023, 33615 Pessac Cedex.

## Abstract

One of the most widespread consequences of climate change is the disruption of trees’ phenological cycles. The extent to which tree phenology varies with local climate is largely genetically determined, and while a combination of temperature and photoperiodic cues are typically found to trigger bud burst (BB) in spring, it has proven harder to identify the main cues driving leaf senescence (LS) in autumn. We used 925 individual field-observations of BB and LS from six *Fagus sylvatica* provenances, covering the range of environmental conditions found across the species distribution, to: (i) estimate the dates of BB and LS of these provenances; (ii) assess the main drivers of LS; and (iii) predict the likely variation in the growing season length (GSL; defined by BB and LS timing) across populations under current and future climate scenarios. To this end, we first calibrated linear mixed-effects models for LS as a function of temperature, insolation and BB date. Secondly, we calculated the GSL for each provenance as the number of days between BB and LS. We found that: i) there were larger differences among provenances in the date of LS than in the date of BB; ii) the temperature through September, October and November was the main determinant of LS in beech, although covariation of temperature with daily insolation and precipitation-related variables suggests that all three variables may affect LS timing; and iii) GSL was predicted to increase in northern beech provenances and to shrink in populations from the core and the southern range under climate change. Consequently, the large differences in GSL across beech range in the present climate are likely to decrease under future climates where rising temperatures will alter the relationship between BB and LS, with northern populations increasing productivity by extending their growing season to take advantage of warmer conditions.

## 1 Introduction

Plants are changing their phenological cycles in response to current climate change (Chmura et al. 2018). Generally, these changes involve a combination of advances in spring leaf phenology and delays in autumn leaf phenology (Gallinat et al. 2015; Piao et al. 2015; Yang et al. 2017), resulting in a longer growing season (Walther et al. 2002; Estiarte and Peñuelas 2015) and potentially increasing forest net ecosystem productivity (NEP) (Way and Montgomery 2015). Phenological responses to environmental cues are to a large extent genetically determined in trees (Liang 2019). Numerous studies along elevational gradients and experiments in common-garden have found bud burst (BB) in populations of different origin to occur at different dates in many tree species (Vitasse et al. 2013; Dantec et al. 2015; Sampaio et al. 2016; Kramer et al. 2017; Cooper et al. 2018). Leaf senescence (LS) has been less widely studied in such settings, but it also differs inherently among populations of *Betula pubescens* (Pudas et al. 2008), *Fraxinus americana* (Liang 2015), *Populus balsamifera* (Soolanayakanahally et al. 2013), *Populus deltoides* (Friedman et al. 2011), *Populus tremula* (Michelson et al. 2018; Wang et al. 2018) and *Populus trichocarpa* (Porth et al. 2015). However, it is not yet clear to what extent the genetic determinism and the environmental cues of BB match those in LS, and how the interplay of BB and LS drives among-population variation in growing-season length (GSL) (Signarbieux et al. 2017).

Extensive research has identified cold winter temperatures (i.e., chilling requirements) and accumulated spring temperatures (i.e., forcing requirements) as the main drivers of BB; sometimes coupled with photoperiod (Basler and Körner 2014; Fu et al. 2015) (Fig. 1). The major drivers of LS have been more difficult to identify (Gallinat et al. 2015; Brelsford et al. 2019). A recent meta-analysis showed that summer and autumn temperatures, precipitation and photoperiod can all affect LS (Gill et al. 2015). Generally, temperature tends to be predominant at lower latitudes (Pudas et al. 2008; Lang et al. 2019), whereas photoperiod is more important at higher latitudes (Soolanayakanahally et al. 2013; Lang et al. 2019) (Fig. 1). Yet temperature effects on LS are not straightforward: increasing summer and autumn temperatures and even moderate drought can delay LS (Xie et al. 2015), whereas severe drought tends to promote earlier LS (Chen et al. 2015; Estiarte and Peñuelas 2015), (Fig. 1). Finally, high insolation may also delay LS (Liu et al. 2016a) (Fig. 1). The complex nature of the environmental triggers of LS has to-date hampered attempts to understand the causes of its variation across large geographical scales (Chmura et al. 2018). This uncertainty makes it very difficult to estimate GSL across species ranges. Recent studies based on *in-situ* records and satellite data have shown positive correlations between the timing of BB and LS that tend to stabilize GSL across populations (Keenan and Richardson 2015; Liu et al. 2016b). But this is not a universal finding and the extent to which GSL can change depends on the combination of many factors, as explained in Fig. 1.

**Figure 1.**
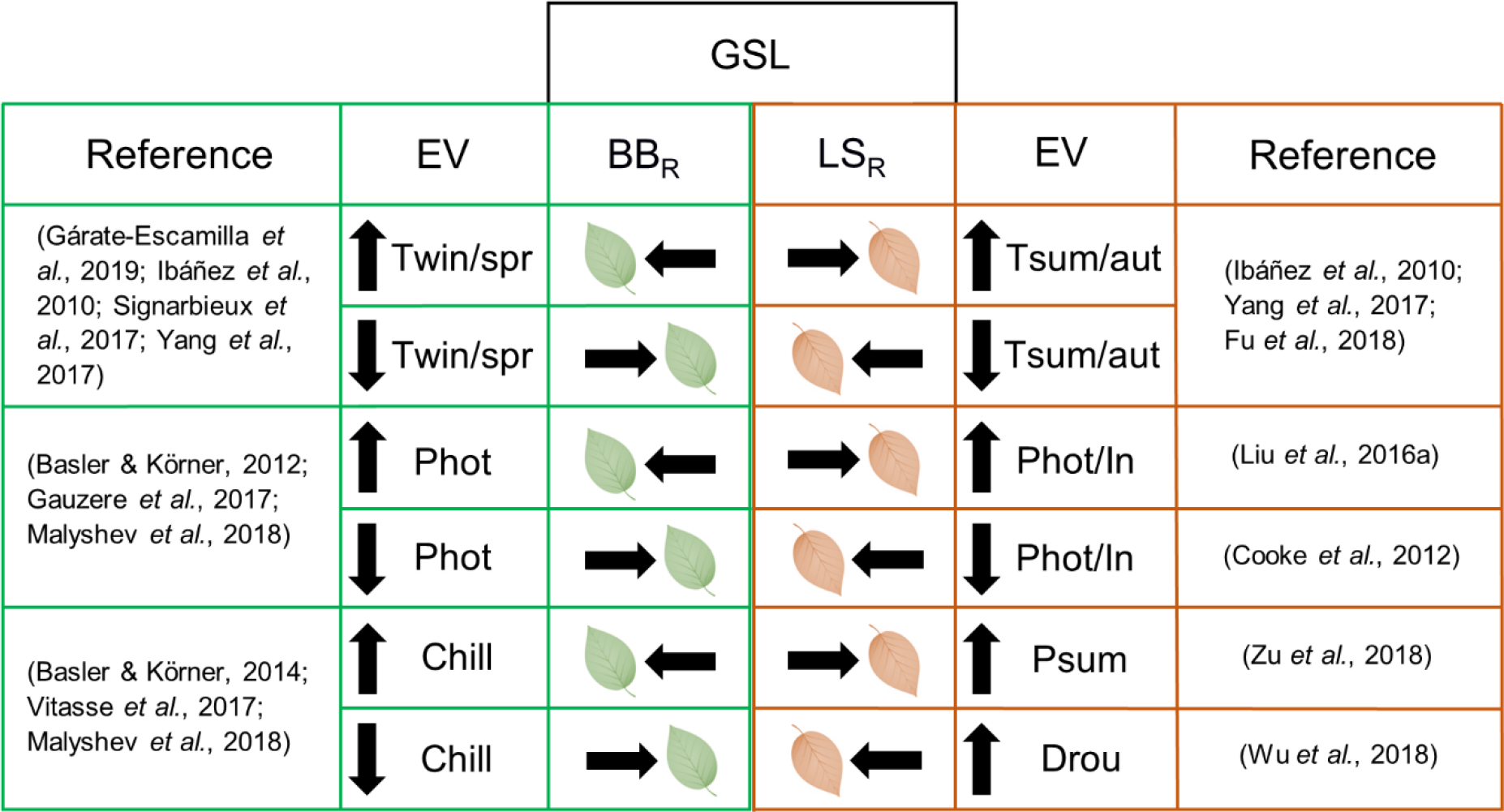
Environmental drivers of growing season length through their effects on bud burst and leaf senescence. GSL: growing season length; EV: environmental variables; BBR: bud burst response; LSR: leaf senescence response; Twin/spr: winter and spring temperatures; Tsum/aut: summer and autumn temperatures; Phot: photoperiod; In: insolation; Chill: chilling requirements; Psum: summer precipitation; Drou: drought; Columns EV: up arrow: increase in the environmental variable; down arrow: decrease in the environmental variable; Columns BBR and LSR: left arrow: early bud burst/leaf senescence; right arrow: delayed bud burst/leaf senescence; Green color and green leaf: Ref, EV related to bud burst and BBR; Orange color and orange leaf: Reference, EV related to leaf senescence and LSR. All the combinations of bud burst and leaf senescence responses defining the growing season length are possible.

*Fagus sylvatica* L. (European beech, henceforth “beech”) is one of the most dominant and widespread broadleaf forest trees in Europe (Preston and Hill 1997) of high ecological and economic importance (Packham et al. 2012). In beech, BB responds to a combination of chilling and forcing temperature requirements (Heide 1993; Falusi and Calamassi 2012; Kramer et al. 2017), and to photoperiod (Heide 1993; Caffarra and Donnelly 2011; Basler and Körner 2012), with the strength of these drivers changing along environmental gradients. For instance, BB is more affected by photoperiod in colder populations, and by chilling requirements in warmer populations (Gárate-Escamilla et al. 2019). Studies of LS in beech suggest that: (i) temperature may be a more important cue than photoperiod when nutrients and water are not limiting (Fu et al. 2018); (ii) non-senescent green leaves are prematurely lost as a result of severe drought conditions (Bréda et al. 2006); (iv) early BB correlates with early LS (Fu et al. 2014; Chen et al. 2018; Zohner et al. 2018); (v) leaves first start to change color in autumn from the upper part of the canopy, suggesting that hydraulic conductance or the amount of solar radiation received over the growing season may play a role in triggering LS (Gressler et al. 2015; Lukasová et al. 2019), although this could also be related to an hormonal effect (Zhang et al. 2011).

Here, we investigate BB and LS in six different beech populations (925 trees) planted in two common gardens in central Europe (Robson et al. 2018), and use this information to infer how range-wide patterns of beech GSL might evolve under future climate warming. Specifically, we attempt to: (i) estimate the dates of BB and LS, and how they differ among populations; (ii) assess the main drivers of LS; and (iii) predict GSL and how it would vary across populations under current and future climate.

## 2 Materials and Methods

### 2.1 Field trials and provenances

Spring and autumn phenological observations came from two common-garden field-trials (hereafter “trials”), located in Schädtbek (54.30°N, 10.28°E), Germany and Tále, Mláčik, Slovakia (48.62°N, 18.98°E) (henceforth termed “Germany” and “Slovakia” trials, respectively). These trials were planted with seeds collected from 38 populations (hereafter referred to as “provenances”) that roughly cover the entire range of beech (Fig. 2). Seeds were germinated in greenhouses and planted in the trials when two years old, in 1995 (Germany) and 1998 (Slovakia) (details given in Robson et al. 2018). To maintain a balanced design, we used only six provenances from each of the two trials (Supplementary Table S1, Fig. 2), chosen in pairs based on their similar climatic origin (Pearson correlation *r≥*0.98). The provenances were ranked from colder (1) to warmer (6) origins (Fig. 2). Trees growing in Germany were measured at an age of 12 and 13 years old, those in Slovakia at 11 and 12 years old (Table S1).

**Figure 2.**
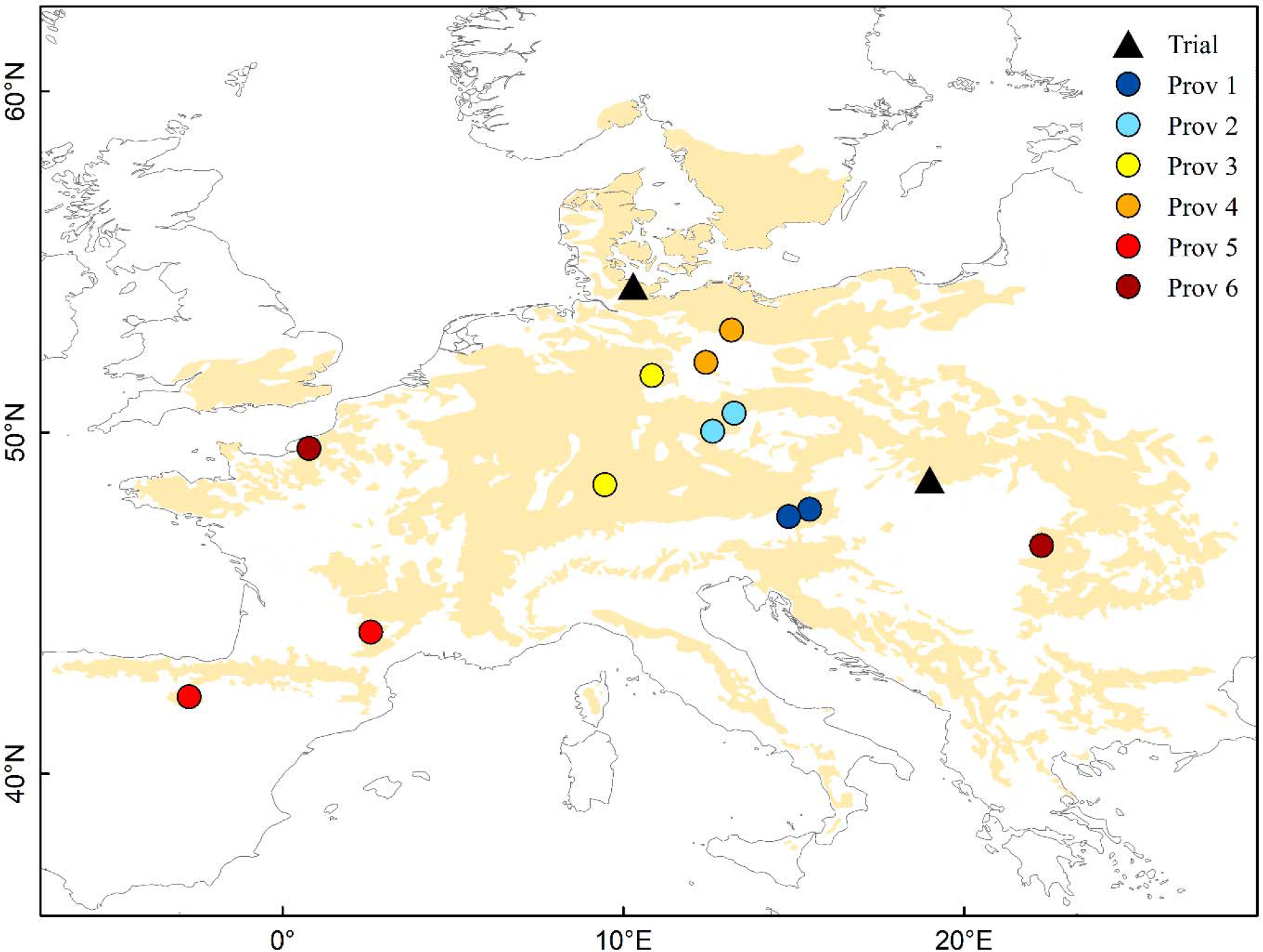
Geographical distribution of beech provenances (colored circles) and trials (triangles) underlying this study. Beige shading indicates the distribution range of beech. The different colors of the provenance circles indicate the pairs of similar provenances selected from each trail (blue colors indicate cold and red colors warm provenances as defined in Table S1).

### 2.2 Estimation of bud burst, leaf senescence and growing season length

We transformed the observational stages (phenophases), and score data (qualitative measurements) for BB and LS to Julian days by fitting the phenophases (Fig. 3; Table S2 and S3) for each tree in every trial using the Weibull function (Robson et al. 2011; Gárate-Escamilla et al. 2019). These data were used to obtain the day of the year (DOY) when BB was attained in spring (stage 2.5; Fig. 2; Robson et al. 2013) and at the stage at which 50% of the trees’ leaves had changed color from green to yellow (stage 3; Fig. 2; (Lang et al. 2019)). We calculated the GSL for each tree as the number of days between the estimated dates of BB and LS (Estiarte and Peñuelas 2015).

**Figure 3.**
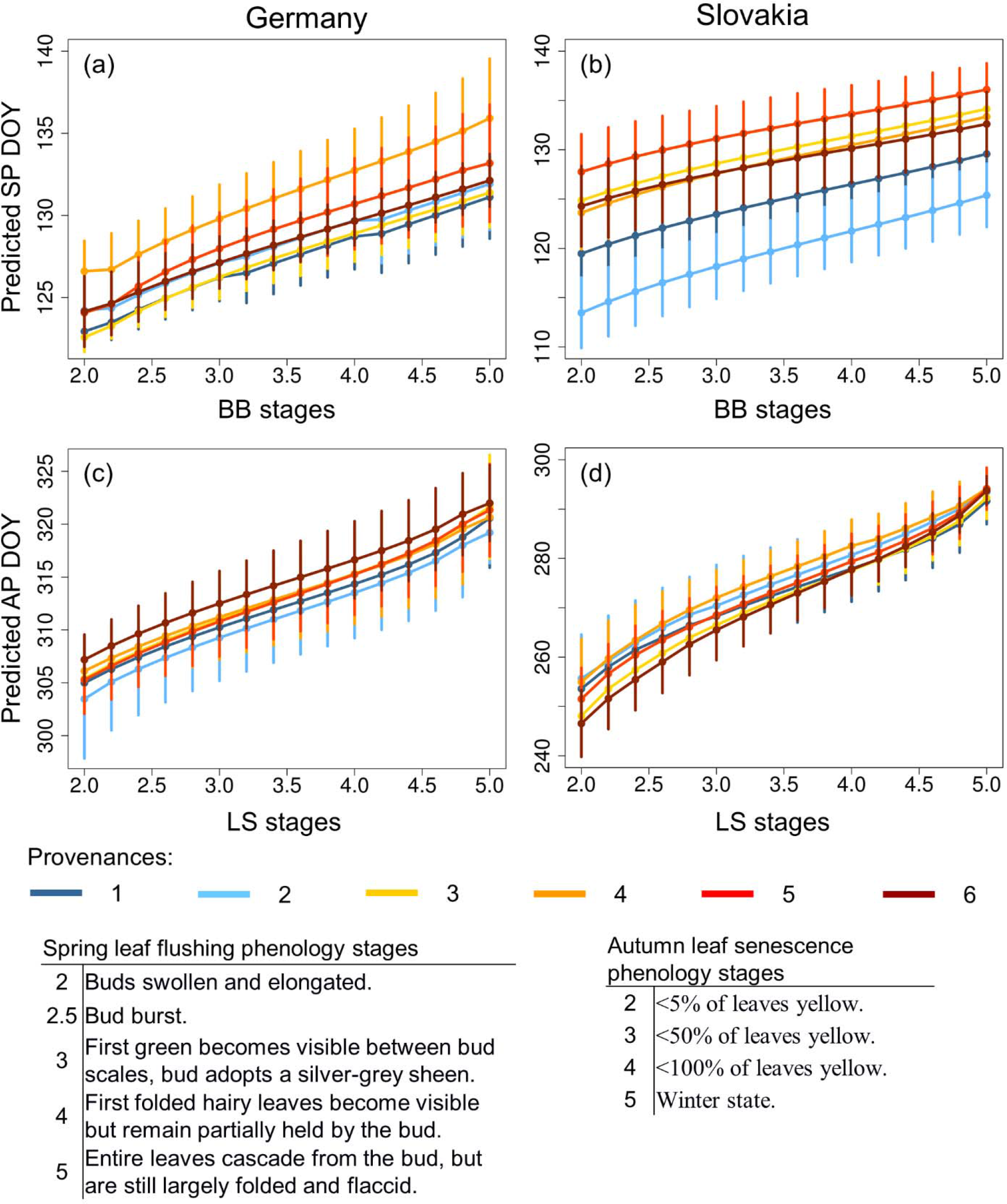
Predicted spring bud burst and autumn leaf senescence phenology days of the year (DOY) against the observational stages recorded in the field for the two trials. SP: spring bud burst phenology; AP: autumn leaf senescence phenology. Provenance colors range from dark blue (cold origin) to dark red (warm origin) for the provenances in the two trials (Figure 1 & Table S1). The spring leaf flushing and autumn leaf senescence stages are described in the lower part of the figure. The phenology stages were recorded in the year 2006 in Germany and 2008 in Slovakia.

### 2.3 Environmental data

To separate the effects of the provenance (genetic effects) to those of the trial (environmental effects), we used the average climate from 1901 to 1990 for each provenance and the average climate of the period between the planting year and the year of measurement for the trial (Leites et al. 2012) in our models. We used the following precipitation- and temperature-related variables from EuMedClim (Fréjaville and Benito Garzón 2018): precipitation of driest month, (BIO14, mm), precipitation (P, mm) of June, July and August (JJA), minimal (Min) monthly water balance (PPET, mm), and mean temperature (Tm, °C) of June, July and August (JJA) and September, October and November (SON). In addition, we used daily insolation, a function of day length and solar irradiance (Yeang 2007). We downloaded daily insolation data from the NASA Atmospheric Science Data Center (https://power.larc.nasa.gov/data-access-viewer/), and we calculated solar radiation (direct and diffuse) between 400-2700 nm incoming on a horizontal surface for a given location. We calculated the mean daily insolation (DIM, kWh m^−2^ d^−1^) between the months of June, July and August (JJA) and September, October and November (SON). As with the climatic variables, we characterized the DIM of the trial as the average between the planting year and the year of measurement. Because the insolation data series from the NASA Atmospheric Science Data Center begins in July 1983, we characterized the DIM of the provenance as the average between 1984 and 1990 for JJA, and between 1983 and 1990 for SON.

We used the 2070 Representative Concentration Pathway (RCP) 8.5 GISS-E2-R (http://www.worldclim.org/cmip5_30s) scenario for GSL predictions under future climate. We deliberately chose only this pessimistic scenario because, for long-lived organisms such as forest trees, it makes little difference whether the projected situation will be reached in 2070 or some decades later.

### 2.4 Statistical analysis

We used a model of BB already calibrated for the same set of trials and provenances (Gárate Escamilla et al. 2019). We then performed a linear mixed-effects model for LS as a function of the combination of environmental variables with BB date as a co-variate. Environmental variables were selected individually to account for separate trial and provenance effects. Our model allowed us to: (i) estimate the date of LS for each of the six pairs of provenances; (ii) compare the date of LS with the date of BB that was already modelled following a similar methodology (Garate Escamilla et al. 2019); (iii) calculate GSL for each provenance; and (iv) perform spatial predictions of BB, LS and GSL under current and future climate scenarios.

#### 2.4.1 Environmental variable selection

To avoid co-linearity and reduce the number of variables to test in our models, we only retained weakly correlated variables (−0.5 < *r* < 0.5) for modeling purposes. The full correlation matrix between all variables is provided in Fig. S2.

#### 2.4.2 Linear mixed-effects model of leaf senescence

We performed a series of linear mixed-effects models of LS as a function of environmental variables from the trial and the provenances, with BB as a co-variable (Equation 1). Each model included one environmental variable from the provenance, one environmental variable from the trial site and BB as fixed effects. The trial, blocks nested within the trial, individual trees and provenances were included as random effects; to control for differences among sites and for repeated measurements of the same tree. The general form of the LS model was:

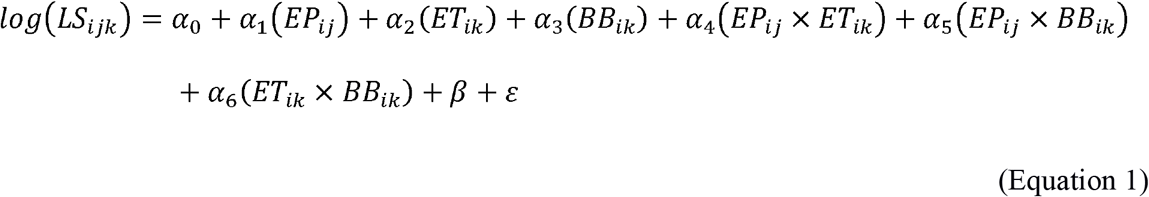

Where LS = leaf senescence of the *i^th^* individual of the *j^th^* provenance in the *k^th^* trial; EP = environmental variable that characterized the provenance site of the *i^th^* individual of the *j^th^* provenance; ET = environmental variable that characterized the trial site of the *i^th^* individual in the *k^th^* trial; BB = bud burst of the *i^th^* individual in the *k^th^* trial; β = random effects and ε =residuals. In addition, the model included the following interaction terms: EP × ET, EP × BB, and ET × BB. EP × ET, interactions represent differences in LS values that can be attributed to the interactions between genetic (provenance) and environmental (site) effects. EP × BB and ET × BB interactions represent the effects of the provenance on LS related to BB and the effects of the site related to BB.

LS models were fitted with the ‘lmer’ function of the package ‘lme4’(Bates et al. 2018), within R statistical framework version 3.2.0 (R Development Core Team 2015). To choose the best supported model, we followed a stepwise-model procedure: (i) we selected the most important variable related to the trial by comparing a series of models that included one environmental variable for the trial and BB, and then selected the best model using the Akaike information criterion (AIC) with criteria delta < 2 (Mazerolle 2006), and the variance explained by the fixed effects, marginal *R*^2^ (Supplementary Table S4); (ii) we chose the optimal random component of the model by comparing the battery of models that included different combinations of random effects, the previously selected environmental variable from the trial and BB using restricted maximum likelihood (REML), and selected using the AIC criterion; (iii) we retained the best environmental variable related to the provenance comparing the models that included one environmental variable from the provenance, the selected variable from the trial, the BB, the interaction between the three variables and the random terms using maximum likelihood (ML) using the AIC criterion (Supplementary Table S4); (iv) we combined the best optimal random and fixed components (previously selected) and adjusted them using REML to obtain the best performing model.

The goodness of fit of the final models was assessed using two approaches. First, we quantified the percentage variance explained by the model attributed to the fixed effects (marginal *R*^2^) and attributed to the fixed and random effects (conditional *R*^2^). Second, we measured the generalization capacity of the model using cross-validation with independent data. To this end, we calibrated the model with 66% of the data and performed an independent validation (using Pearson correlations) with the remaining 34% of the data.

#### 2.4.3 Interactions of leaf senescence with bud burst, and environmental variables

For the best LS supported model, we analyzed the significant interactions (EP × ET, EP × BB, ET × BB in Equation 1) between LS and the environment (ET; represented by one environmental variable of the trial) and according to provenances showing early, mean and late BB. We also inspected gradients of GSL for the six populations by plotting GSL against the environmental variable of the trial selected in the model (ET) and population under current conditions. We predicted the date of LS for the future climate scenario RCP 8.5 using our LS model and the date of BB for the same provenances, achieved using our BB model (Gárate-Escamilla et al. 2019), and plotted the predicted future GSL against ET, for each of the populations.

#### 2.4.4 Spatial predictions

Spatial projections of LS were calculated using our LS model, and predictions of GSL were calculated by subtracting the predicted BB from LS for both current and future climatic conditions across the species range. All spatial predictions were delimited within the distribution range of the species (EUFORGEN 2009). Spatial analyses were performed with the ‘raster’ package in R (Hijmans et al. 2017).

## 3 Results

### 3.1 Estimation of bud burst and autumn leaf senescence dates from field observations

In both trials, differences among provenances were larger for spring leaf flush stages (including bud burst; Fig. 3a & b) than for autumn leaf senescence stages (including 50% yellow leaves; Fig. 3c & d). Although these differences were always statistically significant, they were bigger in the Slovakian trial than in the German one (Fig. 3, Table S2 and S3). Differences in the predicted DOY of spring leaf flush and autumn leaf senescence stages were found for the two different years of measurement in the Slovakian trial (Figs. 3b & d; S1a &b). We used the fitted data to extract the DOY for the flushing stage 2.5 (bud burst, BB) and the senescence stage 3 (= 50% of leaves yellow, LS) for each provenance (Tables S2 and S3).

### 3.2 Variable selection and best model selection

Our inspection of climate variables revealed that: (i) provenance and trial variables were not correlated with each other; (ii) temperature (TmJJA and TmSON)- and precipitation (BIO14, PpetMin and PrecJJA)-related variables for the provenances were correlated, whilst daily insolation (DIMJJA and DIMSON) variables for the provenances were only correlated with the latitude (Lat) of the provenances; (iii) all the trial variables were correlated among themselves; and (iv) the BB co-variable was not correlated with the rest of variables (Fig. S2).

In view of these results, we retained daily insolation (DIMJJA and DIMSON) and temperature-(TmJJA and TmSON)-related variables for the provenances, all climate variables from the trials, and BB as predictors for our models of LS. The best model according to AIC criteria (Tables S4 and S5) used the mean temperature in September, October and November (Tm SON) of the trial and of the provenance, and BB as a co-variable (Table 1 and Table S4).

### 3.3 Leaf senescence model

LS differed among the provenances and between the two trials. These differences were explained by the Tm SON of the trial and provenance, as well as BB (Table 1). Interactions between BB and Tm SON of the trial and provenance were also significant (Table 1). Late BB timing was related to higher Tm SON of the trial and provenances (Fig. 4). Late LS was related to late BB at high Tm SON of the trial, whilst at low trial TM SON the opposite effect occurred (Fig. 4a). Late LS was related to early BB irrespective of Tm SON of the provenance (Fig. 4b). The marginal *R*^2^ was 52%, while the conditional *R*^2^ was 99% (Table 1). The capacity for generalization from the model was *r* = 0.92 (Table 1).

**Table 1.** Statistics from linear mixed-effects models of leaf senescence. Obs: number of trait measurements; Variance: variance explained by the random effects; SD: standard deviation of each level of random effects; Estimate: coefficient of the regression, shown on a logarithmic scale; SE: standard error of each fixed variable; *t*: Wald statistical test that measures the point estimate divided by the estimate of its SE, assuming a Gaussian distribution of observations conditional on fixed and random effects. Fixed effects: Coefficients of the fixed effects of the model; BB: bud burst; Tm SON_T: mean temperature of September, October and November of the trial; Tm SON_P: mean temperature of September, October and November of the provenance. Coefficients of the interactions: BB x Tm SON_T and BB x Tm SON_P. *r*: Pearson correlation; *R*^2^M: percentage of the variance explained by the fixed effects (Marginal variance); *R*^2^C: percentage of the variance explained by the random and fixed effects (Conditional variance).

| Model | Leaf senescence |  |  |
| --- | --- | --- | --- |
|  | Linear Mixed Effect |  |  |
|  | Random Effects |  |  |
|  | Obs | Variance | SD |
| Provenance | 12 | 3.33E-05 | 5.77E-03 |
| Trial | 2 | 2.39E-02 | 1.55E-01 |
| Trial:Block | 6 | 9.73E-06 | 3.10E-03 |
| Tree | 925 | 1.88E-04 | 1.37E-02 |
| Residuals |  | 2.34E-04 | 1.53E-02 |
|  | Fixed Effects |  |  |
|  | Estimate | SE | *t* |
| Intercept | 5.62E+00 | 1.10E-01 | 51.16 |
| BB | -8.18E-04 | 9.91E-05 | -8.25 |
| Tm SON_T | 2.88E-02 | 1.43E-02 | 2.02 |
| Tm SON_P | 2.61E-02 | 8.10E-03 | 3.23 |
| BB x Tm SON_T | 5.97E-04 | 9.61E-05 | 6.21 |
| BB x Tm SON_P | -1.96E-04 | 6.60E-05 | -2.97 |
|  | *r* | *R*<sup>2</sup>M | *R*<sup>2</sup>C |
|  | 0.92 | 0.52 | 0.99 |

**Figure 4.**
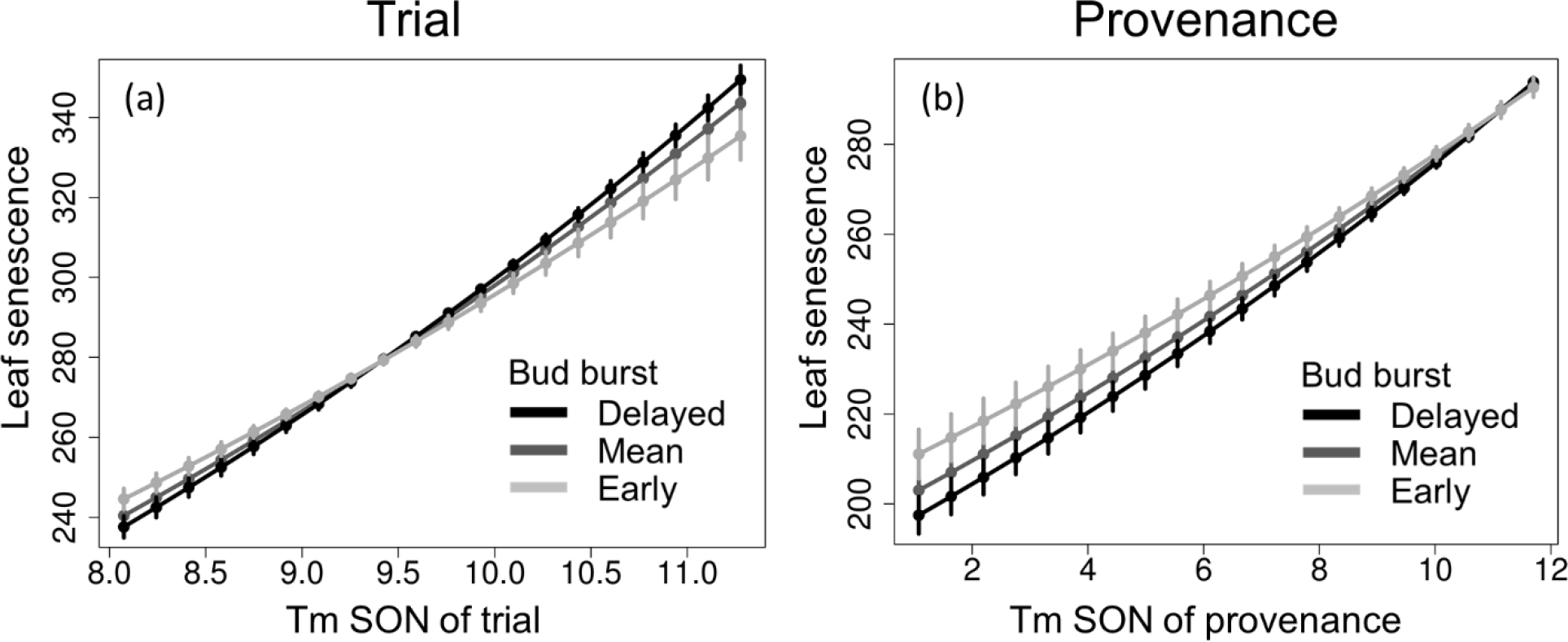
Interaction between leaf senescence and the mean temperature in September, October and November (Tm SON) in the (a) trial and for the (b) provenance. Leaf senescence is given in Julian days, and Tm SON in °C. The black line represents delayed bud burst, the dark gray mean bud burst dates and the light gray early bud burst. The error bars represent the 95% confidence intervals.

### 3.4 Determinants of growing season length under current and future climates

GSL greatly increased with higher temperatures in September, October and November in the trials, although the strength of this effect depended on the origin of the provenances (Fig. 5). Specifically, this increase in GSL was greatest for cold provenances (3.2-5.2 C°), which have their longest GSL under cold conditions (7.5-8.5 C°) at the trials in the current climate (Fig. 5a). In this two specific trials, GSL differed more among provenances under future than under current autumn temperatures (Fig. 5b). The longest GSL under future conditions was predicted at high trial temperatures (11.5-12 C°) for the warm (10.5-11.3 C°) and cold (3.2-5.2 C°) provenances, whilst at low trial temperatures (10.5-11 C°), the longest GSL was predicted for warmer (10.5-11.3 C°) provenances (Fig. 5b).

**Figure 5.**
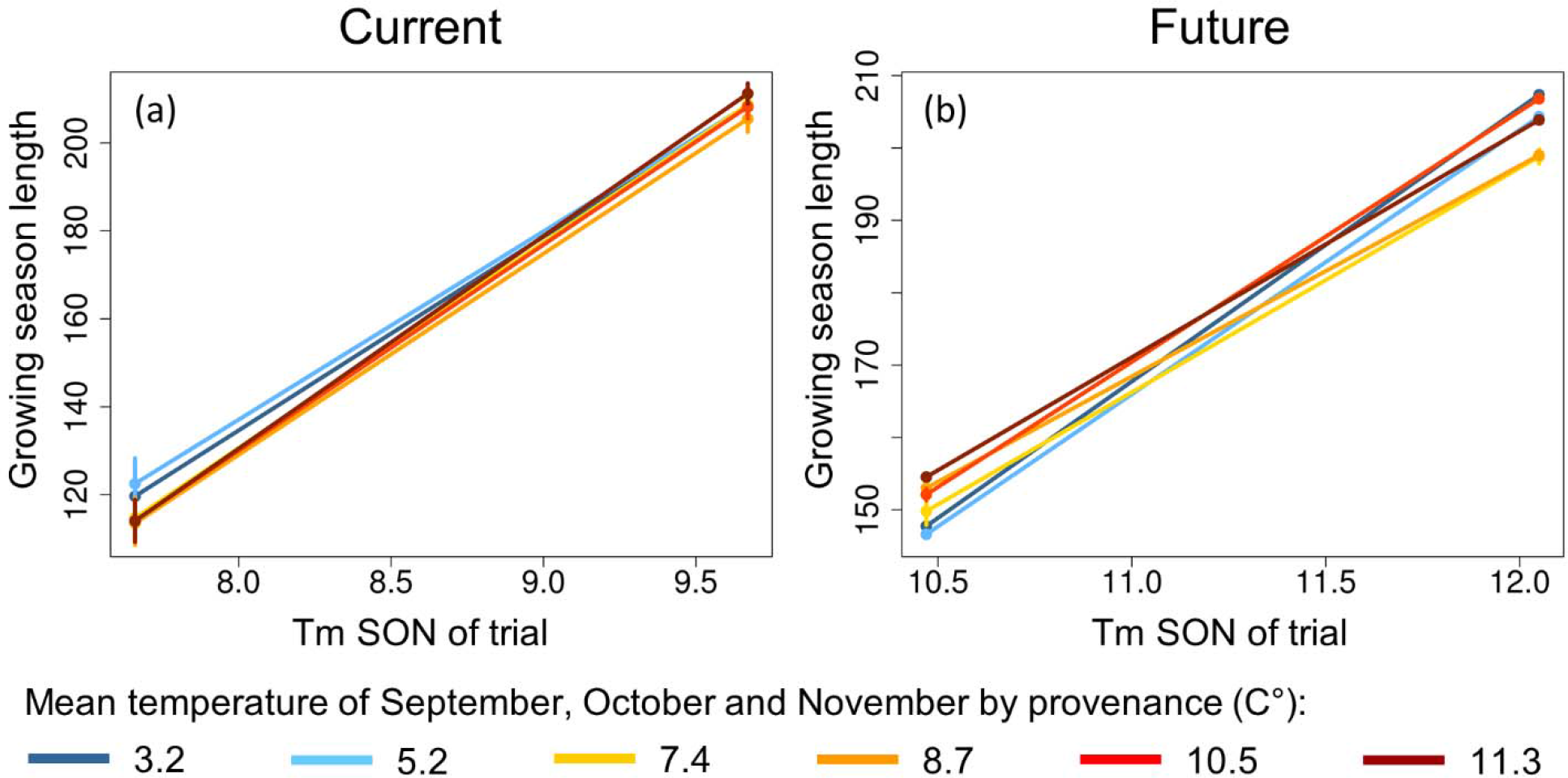
Interaction between growing season length and the mean temperature of September, October and November (Tm SON) of the trial, for (a) current climatic conditions (year of measurement minus year of plantation) and (b) the future climate scenario (RCP 8.5 for 2070). The color gradient depicts the clinal variation from cold (blue) to warm (red) provenances (Tm SON). Growing season length is represented in days. The error bars represent the 95% confidence intervals.

When we extrapolate our models for the examined 2070 climate scenario, GSL is predicted to increase up to 9 days in the northern-east of the range (Fig. 6). Decreases of GSL up to 8 days are predicted for much of the range including the central, southern, western and eastern areas; little or no change in GSL is predicted for the south-eastern-most range (Fig. 6).

**Figure 6.**
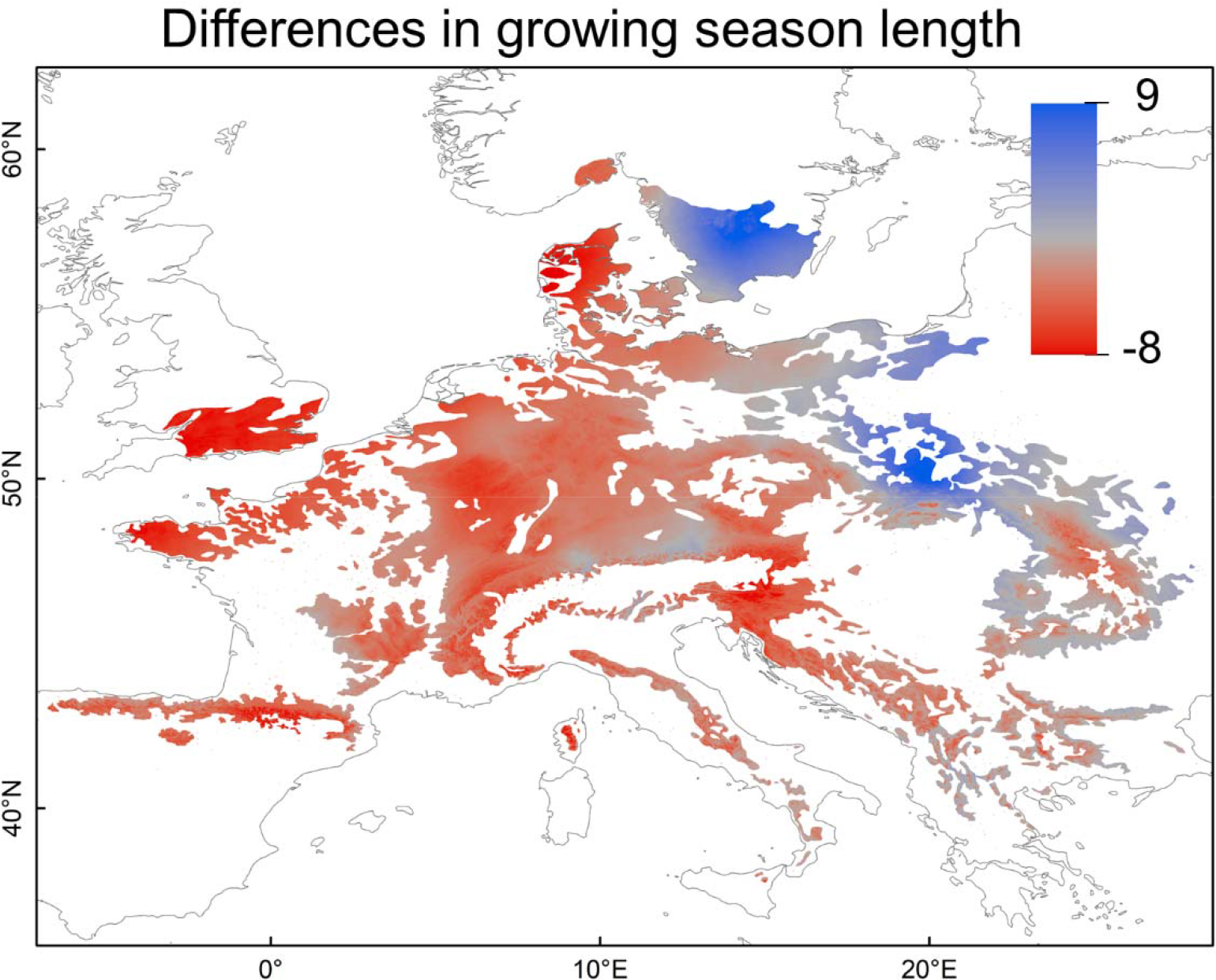
Spatial projections for growing season length: differences between current and future climate conditions. Growing season length is the difference between leaf flushing and leaf senescence spatial predictions (Figure S4). The color gradient depicts the number of days difference in growing season length between current (average climate calculated from 2000-2014) and future conditions (2070, RCP 8.5) from strong decrease (red) to strong increase (blue).

## 4 Discussion

### 4.1 Provenance differences in bud burst and autumn leaf senescence

The origin of beech populations is a major determinant of the timing of their leaf spring and leaf autumn phenology (Table 1), which confirms their genetic differentiation in the control of phenology (Chmura and Rozkowski 2002; Petkova et al. 2017, Alberto et al. 2013). This differentiation is often stronger for spring phenology than for autumn phenology (Vitasse et al. 2009; Weih 2009; Firmat et al. 2017; Petkova et al. 2017), which is in agreement with what we found in our beech provenances (Fig. 3a & b). The duration of autumn leaf senescence is longer than that of leaf flushing in beech (Fig. 3, Table S2, S3) (Gömöry and Paule 2011; Petkova et al. 2017), whereas other temperate broadleaf species such as *Salix spp*. and *Quercus petraea* have a relatively long period of leaf-out and relatively abrupt autumn leaf senescence (Weih 2009; Firmat et al. 2017). Although the dates of spring and autumn leaf phenological stages varied between the two years of our study, the same response-patterns persisted in both years (Figs. 3 and S1), suggesting a consistent effect of environmental conditions on the trials (Weih 2009; Friedman et al. 2011; Petkova et al. 2017). Our results also revealed larger differences among populations for both BB and LS in the Slovakian trial than in the German one (Fig. 3), confirming that, in addition to genetic effects, the environment plays an important role in the phenological response of beech (Vitasse et al. 2013; Gárate-Escamilla et al. 2019).

### 4.2 Environmental variables defining leaf senescence

Overall, our results support the assertions that (1) high autumn temperatures, of the provenance and at the planting site, delay LS in beech, and (2) early BB tends to be followed by early LS (Table 1). The delayed LS promoted by warmer temperatures, that we obtained by manipulating both genetic and site factors using common-garden trials (Fig. 4), is consistent with previous studies based on *in-situ* LS records (Delpierre et al. 2009; Vitasse et al. 2011), satellite data (Yang et al. 2015; Liu et al. 2016a) and climate-controlled chambers (Gunderson et al. 2012; Fu et al. 2018). While the convergence of these studies is reassuring, the extent to which warmer temperatures promote delayed LS still remains elusive (Estiarte and Peñuelas 2015): warmer temperatures accompanied by moderate drought appear to delay LS until a certain threshold (Xie et al. 2015); but beyond this drought threshold LS is accelerated (Chen et al. 2015; Estiarte and Peñuelas 2015). The roles of temperature and drought in LS have several broader implications because the delay in LS induced by warm temperatures is associated with delayed degradation of chlorophyll (Fracheboud et al. 2009), maintenance of photosynthetic enzyme activity (Shi et al. 2014), prolonged leaf life span (Liu et al. 2018a), reduced potential for autumn frost damage (Hartman et al. 2013) and a possible increase of photosynthetic carbon assimilation related to a longer growing season (Liu et al. 2016b).

The finding of our study does not necessarily imply that LS timing in beech only depends on temperature, because this parameter covaried with daily insolation and precipitation (Fig. S2). Both explained a lower proportion of the overall variance (higher insolation promoting delayed LS and higher precipitation promoting earlier LS; see Table S4), yet we cannot exclude the possibility that they may have affected LS timing to some extent (e.g. in parts of the species range not well captured by our model). For instance, insolation can have a strong effect on LS at high latitudes (Liu et al. 2016b) where increasing photosynthetically active radiation with insolation supports increased photosynthesis (Bonan 2002), allowing a delay in LS as a result of persistent chlorophyll retention under sustained high irradiances (Kim et al. 2008).

### 4.3 The effect of bud burst on leaf senescence

The significant carry-over effect of BB on LS timing that we found is consistent with other recent studies on beech (Fu et al. 2014; Signarbieux et al. 2017; Chen et al. 2018; Zohner and Renner 2019), and other deciduous trees across the northern hemisphere (Keenan and Richardson 2015; Liu et al. 2016b). However, it can be difficult to disentangle the effects of temperature on both BB and LS, from their interdependency. In this respect, the significant interaction-effect of BB and the autumn temperature of the provenances on LS is notable (Table 1; Fig. 4), as it suggests that the relationship between BB and LS is moderated by the temperature at provenance origin in a provenance-specific manner. The relationship between BB and LS is complex and various different mechanisms that have been proposed to explain carry-over effects of BB on LS, according to the particular conditions in each study: (i) leaf structural and morphological traits constrain leaf life span (Reich et al. 1992) and programmed cell death (Lam 2004; Lim et al. 2007); (ii) once a plant’s carbohydrate storage capacities are saturated, growth is inhibited (“sink limitation”) and LS is promoted (Fatichi et al. 2013; Keenan and Richardson 2015; Körner 2015; Signarbieux et al. 2017); (iii) LS is itself affected by the preceding winter/spring temperature (Fu et al. 2014; Signarbieux et al. 2017; Zohner and Renner 2019); (iv) early BB could lead to soil water depletion through increased transpiration and resulting in drought stress, producing earlier LS (Buermann et al. 2013); (v) earlier BB might increase pest attack (Jepsen et al. 2011) and increase the probability of spring frost damage (Hufkens et al. 2012), leading to an earlier LS. Our use of multiple provenances of different climatic origin enabled us to isolate the genetic component of these carry-over effects of BB on LS from the temperature response. We only found this pattern among cold provenances (3.2-5.2 C°) (Fig. S3) and in regions with high autumn temperature (11.5-12 C°) (Fig. 4a). Consequently, of the potential causes of this effect, we can discard effects of the preceding winter/spring temperature and of frost damage, neither of which affected the timing of LS in our analysis (results not shown). Yet, we can not rule out the other mechanisms listed above, and more experimental testing is needed to tease apart the relationship between BB and LF across large environmental gradients.

### 4.4 Variation in growing season length based on bud burst, leaf senescence and the environment under present and future climates

Our results, based on two trials located in the core of the distribution range, predict that almost all the provenances monitored (except number 3 – with an average autumn temperature of 7.4°C) would extend their GSL by up to 10 days under future climatic conditions with increased autumn temperatures (11.5-12 C°) (Fig. 5b). However, when we extrapolate our models to the entire distribution range of the species, only trees in northern and north-eastern regions of the species range are predicted to increase their GSL by up-to 9 days, while the GSL of trees in the rest of the range would decrease by up to 8 days (Fig. 6). While several recent studies based on field or satellite data also predict an increase in GSL (Barnard et al. 2018; Liu et al. 2018b; Gaertner et al. 2019) at the high latitudes coincident with cold beech provenances, there have been no recorded increases in the GSL for southern populations of four temperate European species (*Quercus robur, Fagus sylvatica, Betula pendula* and *Aesculus hippocastanum*) over the last two decades (Chen et al. 2018). These two trends are both reflected in our spatial projection of GSL (Fig. 6). The predicted larger differences in GSL in the central and southern range are mostly due to a later leaf senescence predicted for these regions (Fig. S4), which is likely due to an expected increase in autumn temperatures in these regions. We should however note that our spatial modelling results, despite covering a wide climatic range, should be interpreted with caution since they are based on empirical data from only two trials, which can limit their scope.

## 5 Conclusions

European beech is characterised by extensive plasticity in many of its life history traits (Gárate-Escamilla et al. 2019) compared to other tree species (Benito Garzón et al. 2019). Yet strong genetic control over beech phenology, particularly in spring (Kramer et al. 2017), can constrain the acclimative response of populations to climatic changes and hence potentially compromise their future performance. Our analyses provide important insights into the complex relationships driving spring and autumn phenology across the species range. We found large differences in GSL (as inferred from BB and LS) under present climate conditions that are however likely to decrease in the future, because GSLs of southern and core populations (i.e. those with a relatively long current GSL) are predicted to decrease, whilst those of northern and north-eastern populations (i.e. those with a relatively short current GSL) are predicted to increase. These trends are largely driven by an increase in temperatures that would modify phenology. Taken altogether, our results suggest that northern populations would increase productivity in the coming years, extending their growing season to take advantage of warmer conditions in the northern part of the range.

## Founding

This study was funded by the Investments for the Future programme (IdEx) Bordeaux(ANR◻10◻IDEX◻03◻02). HGE was funded by the Consejo Nacional de Ciencia y Tecnologia(CONACYT◻J Mexico; grant number: 636246) and by the Institute of Innovation and Technology Transfer of Nuevo Leon, Mexico. CCB and TMR were funded by the Academy of Finland (decision 304519).

Declaration of Interest: none.

